# Nigral Transcriptomic Profiles in Engrailed-1 Hemizygous Mouse Models of Parkinson’s Disease Reveal Upregulation of Oxidative Phosphorylation-Related Genes Associated with Resistance to Dopaminergic Neurodegeneration

**DOI:** 10.1101/2023.03.22.533733

**Authors:** Lautaro Belfiori, Alfredo DueñasRey, Dorottya Mária Ralbovszki, Itzia Jimenez-Ferrer, Filip Bäckström, Sagar Shivayogi Balikai, Dag Ahrén, Kajsa Brolin, Maria Swanberg

## Abstract

1.

Engrailed 1 (EN1) is a conserved transcription factor essential for programming, survival, and maintenance of midbrain dopaminergic neurons. *En1*-hemizygosity (*En1*^+/-^) leads to a spontaneous Parkinson’s disease-like (PD-like) progressive nigrostriatal degeneration as well as motor impairment and depressive-like behavior in SwissOF1 (OF1*-En1*^+/-^) mice. This phenotype is absent in C57Bl/6j (C57*-En1*^+/-^) mice. Here we studied PD-like phenotypes and early transcriptome profiles in OF1 wild-type (WT) and OF1-*En1*^+/-^ male mice and compare to that of C57 WT and C57-*En1*^+/-^ male mice. To detect transcriptional changes prior to dopaminergic cell loss, we performed RNA-seq of 1-week old mice substantia nigra pars compacta (SNpc). Histology and stereology were used to assess dopaminergic nigrostriatal pathology in 4 and 16 weeks old mice. OF1-*En1*^+/-^ mice showed an increase (±79%) in dopaminergic striatal axonal swellings from 4 to 16 weeks and a loss (±23%) of dopaminergic neurons in the SNpc at 16 weeks compared to OF1 WT. Axonal swellings were also present in C57-*En1*^+/-^ mice but did not increase over time. 52 differentially expressed genes (DEGs) were observed between the C57-WT and the C57-*En1*^*+/-*^ mice, while 198 DEGs were observed in the OF1 strain. Enrichment analysis revealed that the neuroprotective phenotype of C57-*En1*^+/-^ mice was associated with an upregulation of oxidative phosphorylation-related genes compared to both C57 WT and to OF1-*En1*^+/-^ mice.

These results highlight the importance of considering genetic background in PD models and provide valuable insight on how expression of mitochondrial proteins before the onset of neurodegeneration is associated to vulnerability of nigrostriatal dopaminergic neurons.

**Significance statement:** Most PD cases are idiopathic and caused by a complex interplay between genetic variants and environmental risk factors. However, the underlying mechanisms remain elusive. Here we show that *En1* hemizygosity leads to progressive nigrostriatal degeneration with a loss of dopaminergic neurons in OF1-*En1*^+/-^ but that C57-*En1*^+/-^ mice only exhibit early signs of nigrostriatal pathology and do not progress to a PD-like phenotype over time. We identified differences in gene expression related to oxidative phosphorylation before the onset of neurodegeneration to be associated to the differential susceptibility to *En1*^+/-^ induced PD-like pathology. Our work shows how gene expression changes modulate vulnerability to dopaminergic neurodegeneration in the *En1*^+/-^ mouse and reveals putative molecular mechanisms behind the onset and progression of PD.

## 3. Introduction

Parkinson’s disease (PD) represents the fastest-growing neurodegenerative condition (Dorsey et al., 2018). PD is characterized by the loss of dopaminergic neurons in the substantia nigra pars compacta (SNpc) and intracellular aggregates of *α*-Synuclein (*α*-Syn) in the form of Lewy bodies and neurites. PD symptoms include rigidity, bradykinesia, and tremor, and non-motor symptoms like anosmia, constipation, depression, and sleep disorders. Current treatment options temporarily alleviate motor symptoms; however, they do not target the underlying pathology (Poewe et al., 2017; Bloem et al., 2021). A combination of genetic and pathophysiological evidence suggests mitochondrial malfunction (Błaszczyk, 2018), neuroinflammation (Grotemeyer et al., 2022), impairment of lipid metabolism (Galper et al., 2022), autophagy and dysfunctional unfolded protein response (Costa et al., 2020) to be part of PD etiology and pathology.

The genetic component of PD has been extensively studied in familial cases with disease-causing mutations in e.g., SNCA, LRRK2 and PINK1, and in case-control cohorts by genome-wide association studies (GWAS). A recent meta-analysis identified 90 risk loci that account for 16-36% of the heritable PD risk (Nalls et al., 2014, 2019). There is thus an important, yet unknown, genetic component that influences the incidence and progression of PD (Liu et al., 2021; Tan et al., 2021).

To provide a biological context to interpretate the results generated by GWAS, transcriptomics approaches have been used in both murine and human SNpc (Hook et al., 2018; Dick et al., 2020; Monzón-Sandoval et al., 2020; Zaccaria et al., 2022). These studies provided insight on gene expression changes in the SNpc associated with PD. However, most of the generated data relies on the analysis of samples where the neurodegenerative process is already well advanced. Therefore, we lack biological insight into the processes underlying dopaminergic neurodegeneration.

Genetic animal models of PD have proven to be very useful in recapitulating key features of the disease (Konnova and Swanberg, 2018). One of them is the *Engrailed-1* hemizygous (*En1*^+/-^) mouse model, where one copy of *En1* is disrupted by a *LacZ* insertion (Saueressig et al., 1999). The EN1 protein is a highly conserved homeodomain transcription factor essential for the development and survival of the mesodiencephalic dopaminergic neurons, along with Engrailed-2 (*En2*) and pentraxin-related gene 3 (*Ptix3*) (Veenvliet et al., 2013). EN1 also has a role in the maintenance and sustainment of energy metabolism of dopaminergic neurons in adulthood (Rekaik et al., 2015a). Furthermore, *En1*^+/-^ mice exhibit a spontaneous and thoroughly characterized PD-like neuropathology which recapitulates PD in terms of progressive loss of dopaminergic neurons in the SNpc but sparing of VTA neurons, mitochondrial deficits, axonal degeneration, *α*-Syn-positive aggregates and diminished dopamine release in the dorsal striatum (Sonnier et al., 2007; Alvarez-Fischer et al., 2011a; Fuchs et al., 2012; Nordström et al., 2015; Rekaik et al., 2015b; Chatterjee et al., 2019).

We and others have previously shown that the consequences of *En1* hemizygosity in mice depends on genetic factors (Sgadò et al., 2006a; Murcia et al., 2007; Sonnier et al., 2007) and that these are located outside the En1 and En2 loci (Kurowska et al., 2016). While *En1*^+/-^ SwissOF1 (OF1-*En1*^+/-^) mice display the spontaneous neurodegenerative phenotype, *En1*^+/-^ C57Bl/6j (C57-*En1*^+/-^) mice require an additional deletion of *En2* to develop a neurodegenerative phenotype (Sgadò et al., 2006b). We have found that the protective effect of the C57 background genome is complex, and maps to 23 partly overlapping and interacting quantitative trait loci (QTL) linked to dopaminergic cell loss in SNpc and axonal pathology (Kurowska et al., 2016).

Here, we aimed to identify processes and pathways associated to susceptibility and resistance to PD-like pathology in *En1*^+/-^ mice with different genetic backgrounds. To achieve this, we analyzed transcriptomic profiles of nigral neurons from OF1-*En1*^+/-^ and C57-*En1*^+/-^ mice at very early stages of the neurodegenerative phenotype. Our results provide evidence that increased expression of genes encoding mitochondrial proteins regulating oxidative phosphorylation protect C57-*En1*^+/-^ mice from dopaminergic neurodegeneration, further supporting energy metabolism as an important player and therapeutic target in PD.

## 4. Materials and methods

### 4.1. Animal husbandry, breeding scheme and experimental setup

All the experiments were approved by the local ethics committee Malmö/Lund region and specified if permit 4992/2022. Male mice were housed under a 12-hour Light: Dark cycle with free access to food and water. Wild type (WT) Females were obtained from (Breeder/company and full name of strain) and mated with *En1*^+/-^ males (C57-*En1*^+/-^ and OF1-*En1*^+/-^). OF1-*En1*^+/-^ were generated as described earlier (Sonnier et al., 2007). C57-*En1*^+/-^ mice were generated by repeated backcrossing of OF1-*En1*^+/-^ to C57 background (Van Andel Institute, MI, USA). Mice were sacrificed at 4 different ages: 1, 4 and 16 weeks. Seven mice per genotype and strain were used for histological analysis. As for RNA-Seq, six animals from each strain were used, of which 3 were WT and 3 *En1*^+/-^. Exclusively male mice were used in this study. Four groups of male mice were studied in order to assess both *En1*^+/-^ and genetic background effects (Fig. 1, A)

**Figure 1.**
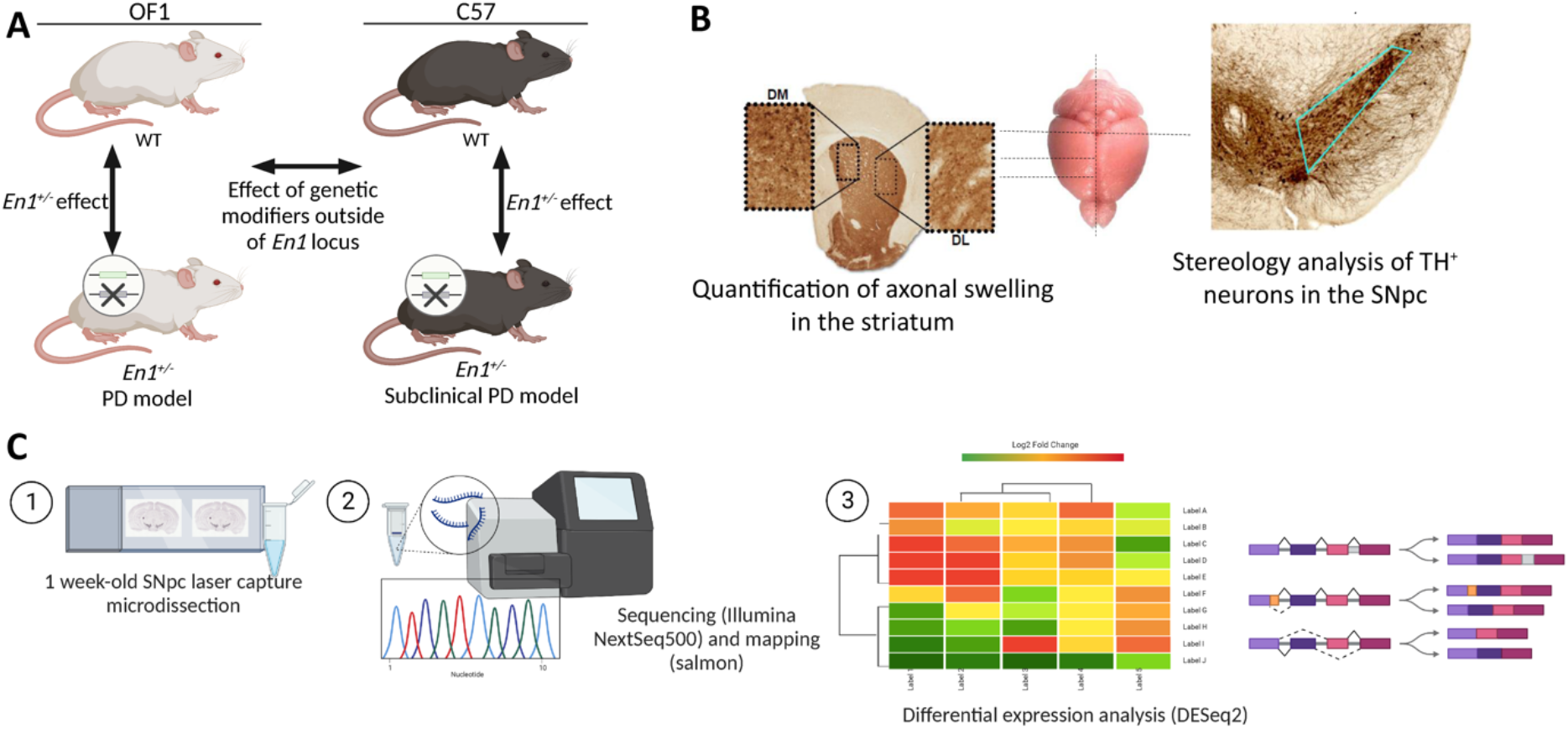
Analysis of genetic background effects on neurodegeneration in the *Engrailed 1* hemizygous (*En1*^*+/-*^) mouse model for Parkinson’s disease (PD). **A**. Study design and experimental setup (n=3). **B**. Sections used for histological analyses of PD-like neuropathology in *En1*^*+/-*^ mice. **C**. Methodology for bulk RNA-Seq analysis of laser capture microdissected substantia nigra pars compacta (SNpc) from 1 week-old mice. OF1: SwissOF1, C57: C57Bl/6J, TH: Tyrosine hydroxylase

### 4.2. DNA isolation and genotyping

Ear or tail punches were used as source of genetic material. KAPA® HotStart mouse genotyping kit (KK7351, Roche, CH) was used to extract genomic DNA and perform the PCR reaction following the manufacturer’s instructions. *En1* hemizygosity was determined using primers specific for *lacZ*, inserted in the *En1* locus, producing a null allele (Supp. Table S1). Amplification conditions were set up as follows: 95°C for 3min, 32 cycles of 95°C for 30s, 58°C for 30s, 72°C for 30s, followed a final extension at 72°C for 7 minutes. To identify the sex of 1 week mice, PCR (94°C for 2min, 32 cycles of 95°C for 30s, 55°C for 30s, 72°C for 30s, followed a final extension at 72°C for 7min) with Y-chromosome specific primers were used (Supp. Table S1). PCR-products were analyzed on 2% agarose gel with SYBR(tm) Safe DNA Gel Stain (X10000, #S33102, Thermo Scientific(tm)).

### 4.3. Immunohistochemistry, axonal swelling quantification and stereology

Four and 16 week old mice were euthanized by intraperitoneal injection of 0.2ml of sodium pentobarbital (60mg/ml), perfused with ice-cold saline (0.9% NaCl) for 5 minutes and fixated by perfusion with 4% paraformaldehyde (PFA, pH7.4) for 5 minutes. Brains were then removed and postfixed in 4% PFA overnight (O.N.). Once fixated, brains were transferred into a 30% sucrose, with 0.01% sodium azide solution in phosphate buffer (PBS) until fully embedded.

For immunohistochemistry (IHC) brains were sectioned coronally on a freezing sledge microtome (Leica SM2010R). Six series of consecutive coronal 40*μ*m thick sections were collected for both striatum and SNpc (Fig 1, B). Sections were stored at 4°C on Walter’s antifreeze (30% (v/v) ethylenglycol, 30% (v/v) glycerol,0.01% (v/v) sodium azide, and 0.5 M phosphate buffer) until tyrosine hydroxylase (TH) staining. Free floating sections were washed 3 times with PBS, quenched for 30 minutes with 3%H_2_O_2_/10% MeOH in PBS. After washing again with PBS, sections were permeabilized by incubating for 10 minutes in 0.3% Triton X-100 in PBS (PBS-T) and blocked with 5% normal goat serum for 2 hours. Next, sections were stained O.N., at 4°C, with TH primary antibody (1:4000, AB152, Millipore). The next day, sections were washed with PBS-T and incubated with a biotinylated secondary antibody (1:200, BA9200, Vector Laboratories, UK) for 1 hour at room temperature. This was followed by washes with PBS-T and PBS, and a 30 min incubation with Avidin-Biotin complex reagent (ABC Elite, Vector Laboratories, UK). Finally, immunostaining was revealed by incubation with diamonebenzidine (DAB) as chromogen. Mounted sections were then dehydrated on increasing concentrations of ethanol, cleared with xylene, and slipcovered using DPX mounting media.

For axonal swelling quantification, three consecutive sections from each animal were used. Pictures were taken at a 20X magnification using an Olympus BX53 microscope (Olympus, Japan). Two pictures per section were taken representing the dorsal part of the caudate putamen. Analysis was done using ImageJ (1.52a, Java 1.8.0_112 64-bits, NIH, USA). Axonal swelling were counted and classified according to their size: small (0.1-1.5*μ*m), medium (1.51-2.99*μ*m) and large (>3*μ*m). Averages were determined based on total pictures for each animal.

Stereology analysis was done following the optical fractionator principle (West and Gundersen, 1990) to estimate the total number of TH^+^ dopaminergic neurons in the SNpc. Analysis was done on every third section of the midbrain of each animal, resulting in 8-10 sections per animal. For imaging and quantification, a Leica MPS52 microscope and the Stereo Investigator® software (MBF bioscience) were used. The parameters include 5X/0.11 lens for delimiting the region of interest (ROI), 100X/1.3 lens for counting, a counting frame size of 55×55*μ*m, a sampling area of 130×130*μ*m, a section thickness of 30.1±0.9*μ*m and a Gundersen coefficient error of ≤ 0.07.

### 4.4. Laser capture microdissection, RNA extraction from nigral tissue, RNA sequencing, and read counting

One week old mice were sacrificed by decapitation; brain dissections were performed on ice under 2 minutes to preserve RNA integrity. Once removed from the skulls, the brains were embedded in Optimal Cutting Temperature Compound (Tissue-Tek® O.C.T. Compound, Sakura® Finetek), snap-frozen in liquid nitrogen, and stored at -80°C until used.

Before the microdissection, Polyethylene naphtholate (PEN)-membrane slides (1mm, 0.17mm; 415190-9041-000, Carl Zeiss MicroImaging, Inc.) and staining jars were heated at 180°C for 4 hours followed by UV light irradiation to completely inactivate RNases and destroy contaminating nucleic acids. Frozen tissue blocks were transferred to a cryostat (CM3050 S, Leica Microsystems) that was pre-set to –17°C and allowed to equilibrate for at least 1 hour before sectioning. A total of six 14µm-thick sections containing early substantia nigra (approximately equivalent to Bregma -2.92 to -3.16 in adult mice) were collected from each experimental animal, accommodating three per slide. Slides were stored at -80°C until staining and Laser Capture Capture Microdissection (LCM), which were performed on the same day. A short staining procedure was conducted and ice-cold solutions were used to preserve the RNA integrity. Briefly, sections were fixed in 70% EtOH for 2min, followed by staining with 1% w/v cresyl violet acetate prepared in 50% EtOH. After removal of excess stain, the slides were dipped in 70% and 100% EtOH, and air-dried before capturing the target region using the PALM Robot Microbeam Laser Microdissection System (Zeiss) at the Division of Oncology and Pathology (Department of Clinical Sciences, Lund University).

AdhesiveCap clear tubes (415190-9211-000, Zeiss) were used for collecting the nigral tissue from each animal separately: approximately 0.3mm^2^ of tissue was harvested from each animal and catapulted onto the adhesive cap. Immediately after LCM, tissue lysis was performed using 200µL RLT buffer with β-Mercaptoethanol from the RNeasy Micro kit (#74004, QIAGEN,) in a ventilated hood for 30min followed by centrifugation at 9000rcf for 5min. Total RNA purification was performed following the RNeasy Micro kit. The quality of the RNA was assessed on an Agilent 2100 Bioanalyzer RNA following the manufacturer’s instructions (G2946-90005, Agilent Technologies, Inc.). Samples were then stored at -80°C until cDNA library preparation and sequencing.

Library construction and sequencing were performed at the MultiPark Next Generation Sequencing facility (Department of Experimental Medical Science, Lund University). Due to the low initial RNA concentrations, the SMART-Seq® v4 Ultra® Low Input RNA Kit (Takara Bio, Inc.) was used for cDNA library preparation; the output was then processed with the Nextera® XT DNA Library Preparation Kit (Illumina, Inc.) for paired-end (2×75bp) sequencing on an Illumina NextSeq500 instrument. Multiplex libraries were sequenced at an average depth of 20 million reads per sample.

After demultiplexing, the quality of the Illumina reads stored in FASTQfiles was assessed by means of FastQC (v0.11.4). Transcript quantification was performed using Salmon (v0.8.2) and the transcriptome-based quasi-mapping model (Patro et al., 2017). The following flags were used for quasi-mapping: *--gcBias* and *--validateMappings*. A Bash script was used to run Salmon on all samples in serial. The transcript index was derived from the GENCODE mouse release M28 (GRCm39) containing nucleotide sequences of all transcripts on the reference chromosomes.

After transcript quantification, the data were imported into the R statistical computing environment (v4.1.2) and summarized using tximport (v1.22) (Fig. 1, C).

### 4.5. Identification of differentially expressed genes (DEGs)

Differential gene expression analysis between (i.e., OF1 vs. C57) and within (i.e., *En1*^+/-^ vs. WT) strains was done using DESeq2 (v1.32) following the authors’ suggested workflow (Love et al., 2014). Transcripts that did not reach a threshold of 10 normalized read counts after averaging across biological replicates were eliminated and considered as not expressed. The model for differential gene expression was parametrized to evaluate strain and genotype effect, and its interactions (∼Genoype+Strain+Genotype:Strain). Contrasts for each of the 4 comparisons (Fig. 1, A) were analysed. Significance for differential expression was accepted at the Benjamini-Hochberg-adjusted p<0.05. Differentially expressed genes (DEGs) with FC| > 1.5 after lfc shrinkage were selected.

### 4.6. Functional enrichment analysis of DEGs

The clusterProfiler (v4.0.5) package (Yu et al., 2012) was used to explore overrepresented biological pathways using the Gene Ontology (GO), Kyoto Encyclopaedia of Genes and Genomes (KEGG) and Wikipathways. GO categories with a Benjamini-Hochberg-corrected p≤0.05 are reported. Reactome pathway-based analysis was done using the ReactomePA package (Yu and He, 2016).

### 4.7. Experimental design and statistical analyses

Statistical tests for the striatal axonal swelling quantification and stereological counts of TH^+^ cells in SNpc of 4- and 16-week-old mice were performed using GraphPad Prism Software (version 7, GraphPad, La Jolla, CA). The Shapiro-Wilk test was performed in R (v4.1.2) to test for normality. Differences between groups were analysed using two-way ANOVA with Tukey’s multiple comparisons test; statistical significance was set at p < 0.05 and values are expressed as mean ± standard deviation (SD) for the histological analyses.

### 4.8. Data availability

The data will be publicly available at GEO repository once the article has been published. Further data can be made available upon reasonable request.

## 5. Results

### 5.1. Both OF1-*En1*^+/-^ and C57-*En1*^+/-^ mice display early nigrostriatal pathology

Previous work has showed signs of nigrostriatal degeneration in OF1-*En1*^+/-^ mice in the form of spheroidal dystrophic terminals (axonal swellings) as early as postnatal day 8, that progressively become abundant at 4 and 16 weeks (Sgadò et al., 2006c; Sonnier et al., 2007; Nordström et al., 2015). We found similar levels of axonal swellings per number of nigral dopaminergic neurons at 4 weeks in OF1-*En1*^+/-^ and C57-*En1*^+/-^ mice (0.047±0.02 vs 0,042±0.01; n=7, p=0.9) (Fig. 2; A, B). While OF1-*En1*^+/-^ mice showed a significant increase in axonal swellings when comparing 4 and 16 weeks (0.047±0.02 vs 0.084±0.02; p=0.0003), axonal pathology did not increase over time in C57-*En1*^+/-^ mice (0.042±0.01 vs. 0.06±0.01, p=0.09). Furthermore, OF1-*En1*^+/-^ had significantly more axonal swellings compared to C57-*En1*^+/-^ mice at 16 weeks (0.084±0.02 vs 0,042±0.01; p=0.02). Two-way ANOVA indicated effects of age (F (1, 24) = 27,21, p<0.0001) and strain (F (1,24) = 7,280, p=0.01), but no interaction between the two factors.

**Figure 2.**
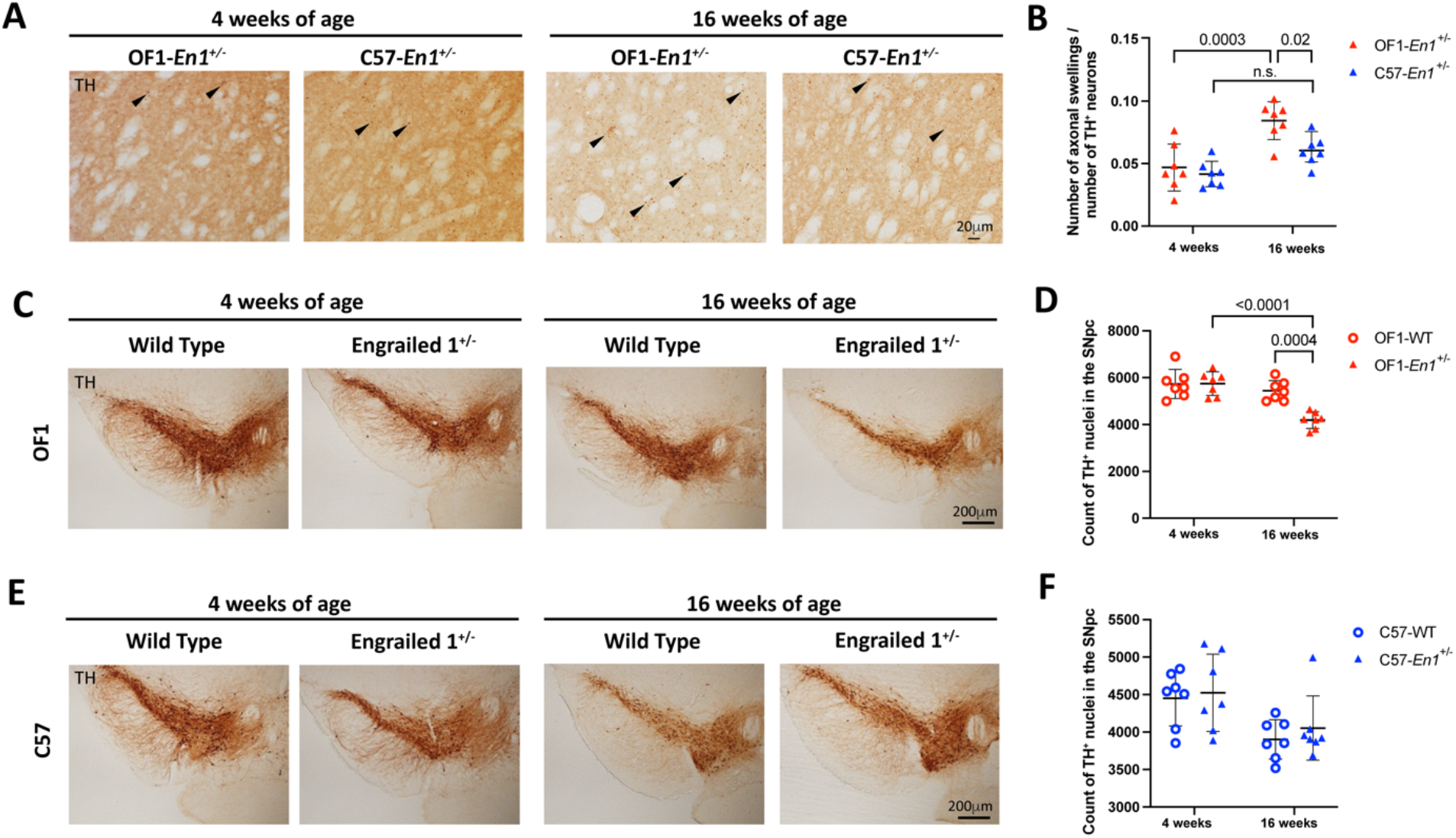
Engrailed 1 hemizygosity (*En1*^*+/-*^) induces progressive dopaminergic neurodegeneration in OF1 but not C57 mice. **A**. Representative images of axonal swellings in the striatum of OF1-*En1*^*+/-*^ and C57-*En1*^*+/-*^ mice at 4 and 16 weeks-of-age. **B**. Axonal swellings are detected in both OF1-*En1*^*+/-*^ and C57-*En1*^*+/-*^ mice at 4 weeks-of-age. In OF1-*En1*^*+/-*^mice, there is a significant increase in the number of swellings per remaining dopaminergic neuron in the substantia nigra pars compacta (SNpc) between 4 and 16 weeks, while axonal swellings do not increase in C57-*En1*^*+/-*^ mice. Data presented as mean ± SD (Two-way ANOVA with Tukey’s multiple comparison test; n = 7). **C**. Representative images of SNpc from OF1 wild type (WT) and OF1-*En1*^*+/-*^ mice at 4 and 16 weeks-of-age. **D**. The number of dopaminergic neurons in the SNpc is significantly lower (∼-23%) in 16 week-old OF1-*En1*^*+/-*^ mice compared to both 16 week-old OF1 WT and 4 week-old OF1-*En1*^*+/-*^ mice. **E**. Representative images of SNpc from C57 WT and C57-*En1*^*+/-*^ mice at 4 and 16 weeks-of-age. **F**. C57-*En1*^*+/-*^ mice did not display any loss of nigral neurons in the SNpc at 16 weeks. The estimated number of dopaminergic neurons in the SNpc of C57-WT mice was lower compared to OF1-WT at both 4 and 16 weeks. TH: Tyrosine hydroxylase, OF1: SwissOF1, C57: C57Bl/6J

Early nigrostriatal pathology is thus manifested in both strains, but while the axonal pathology increases over time in OF1-*En1*^+/-^ mice, it does not progress in C57-*En1*^+/-^ mice. This suggests that genetically determined compensatory mechanisms can counteract a progressive PD-like phenotype.

### 5.2. C57 genetic background protects against dopaminergic neuron loss induced by *En1* hemizygosity

We have previously reported that the loss of dopaminergic neurons due to *En1* hemizygosity is genetically determined by multiple QTLs (Kurowska et al., 2016). To further characterize the nigral degeneration phenotype, we performed stereological counting of tyrosine hydroxylase positive (TH+) neurons in the SNpc of 4 and 16 week mice. OF1-*En1*^+/-^ mice show a ≈23% loss of dopaminergic neurons in the SNpc when compared to OF1 WT mice at 16 weeks (mean dopaminergic neurons number 5442±433 vs 4187±356, n=7, p=0.0004), but no significant cell loss at 4 weeks (Fig.2; C, D). To determine if the phenotype progressed over time, we performed two-ways ANOVA that showed significant effects of age (F (1, 24) = 25.23, p<0.0001), *En1*^+/-^ genotype F (1, 24) = 11.36, p=0.0004) and interaction between the two (F (1, 24) = 11.89, p=0.0021). In contrast, C57-*En1*^+/-^ mice showed no loss of dopaminergic neurons in the SNpc when compared to C57 WT at 4 weeks (4525±514 vs 4451±369) or 16 weeks (4053±429 vs 3902±264) (Fig. 2; E, F). In the C57 mice, two-way ANOVA detected a significant effect of age (F (1,24) = 11.13, p=0.0028), but not for *En1*^+/-^ genotype or an interaction of the. Histology thus confirms a progressive dopaminergic neurodegeneration in OF1-*En1*^+/-^ mice and support that the genetic background in C57-*En1*^+/-^ mice confers resistance to this PD-like phenotype.

### 5.3. Transcriptomic analysis of laser captured micro-dissected SNpc

To assess factors driving or counteracting the neurodegenerative process in its early stages, we performed RNA-sequencing of laser captured micro-dissected (LCM) SNpc from 1 week old OF1 WT, OF1-*En1*^+/-^, C57 WT and C57-*En1*^+/-^ mice (n=3 per group). We achieved precise isolation of SNpc through LCM (Sup.Fig. 1, A) and obtained high quality RNA (RIN > 8.0) (Sup.Fig. 1, B). All samples showed high Phred quality scores (>30) after sequencing, as assessed by FastQC (v0.11.8) software with most of the reads being 75bp long (Sup. Fig. 1C, D).

Initial analysis of the data through principal components analysis (PCA) showed that the main effect was due to genetic background. *En1* hemizygosity induced more differentially expressed genes (DEGs) in the OF1 than C57 strain, while several DEGs were shared between the comparisons evaluating strain differences (Sup. Fig. 2, A, B). A decreased expression of *En1* was seen in both OF1-*En1*^+/-^ and C57-*En1*^+/-^ mice compared to their respective WT, but with low counts, high variation, and no significant differences (Sup. Fig. 2, C).

### 5.4. Distinct transcriptomic profiles are induced by *En1* hemizygosity in OF1 and C57 mice

Firstly, we aimed to study the effects of *En1* hemizygosity on transcriptional profiles in OF1 and C57 mice. Analysis of OF1-*En1*^+/-^ vs OF1 WT and C57-*En1*^+/-^ vs C57 WT detected 198 and 52 DEGs, respectively (FDR < 0.05) (Fig. 3, A). Among the DEGs, 141 were exclusive to OF1 and 36 genes were exclusive to C57. Only two genes (*Pax3* and *Smc2*) were differentially expressed in both OF1-*En1*^+/-^ and C57-*En1*^+/-^ compared to WT mice (Fig. 3, B).

**Figure 3.**
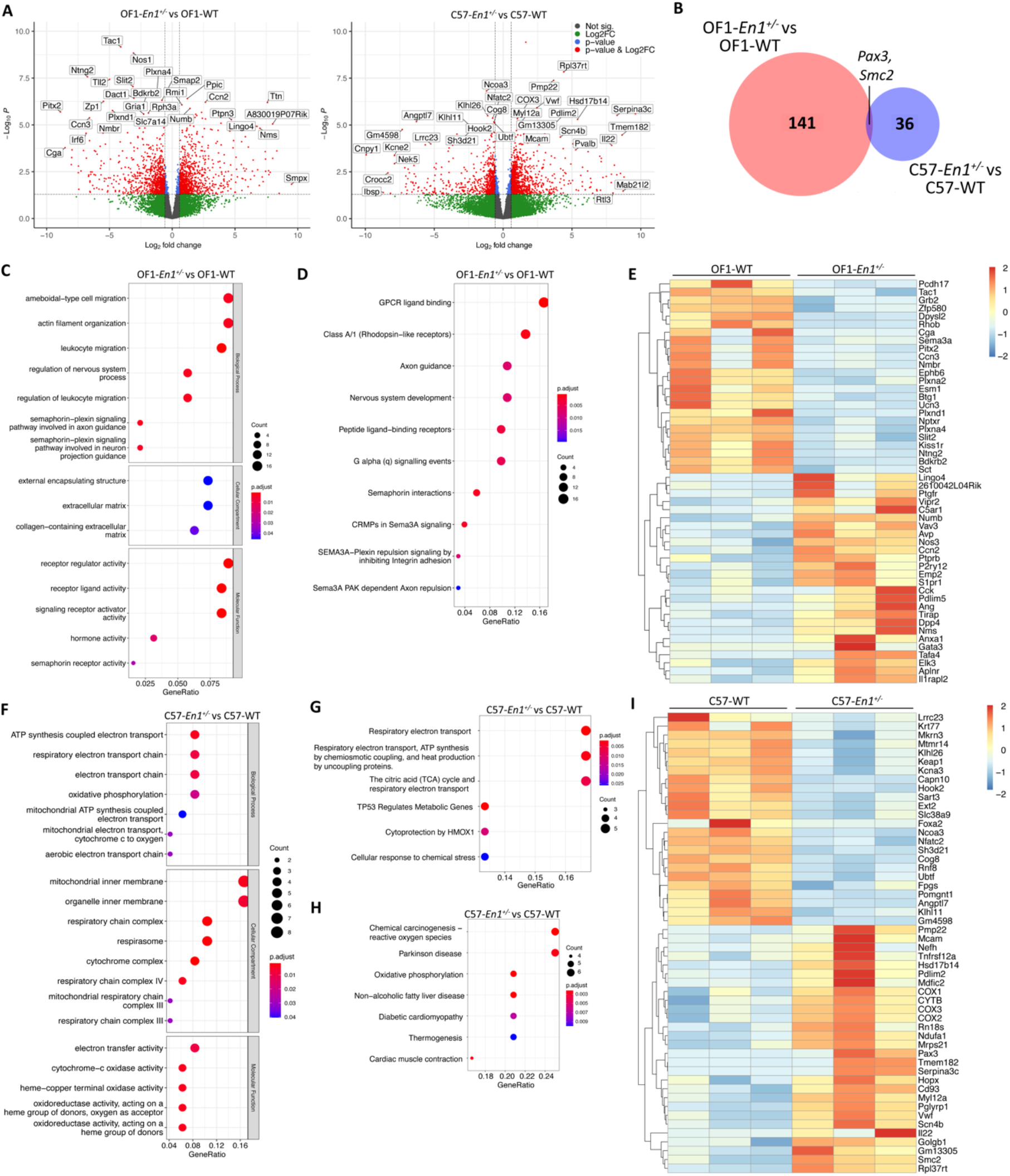
Analysis of Engrailed-1 hemizygosity effects on nigral transcriptomic profiles in OF1 and C57 mice. **A**. Volcano plot showing all expressed genes in comparisons between OF1-*En1*^*+/-*^, C57-*En1*^*+/-*^ and their respective wild type (WT) strain. Vertical lines indicate fold change (|FC|) of 1.5. **B**. Venn diagram illustrating the small overlap between differentially expressed genes (DEGs) detected in OF1*-En1*^*+/-*^ vs. OF1 WT and C57*-En1*^*+/-*^ vs. C57 WT. **C**. Gene Ontology (GO) enrichment analysis of DEGs between OF1-*En1*^*+/-*^ and OF1 WT mice. **D**. Reactome pathway analysis for DEGs between OF1-*En1*^*+/-*^ and OF1 WT mice. **E**. Heatmap of genes included in the enriched pathways among DEGs between OF1-*En1*^*+/-*^ and OF1 WT mice. OF1 WT mice show a lower expression of genes associated with neuron development and axonal guidance. Red: up-regulated, blue: down-regulated. **F**. GO enrichment analysis of DEGs between C57-*En1*^*+/-*^ and C57 WT mice. **G**. Reactome pathway analysis for the DEGs between the C57-*En1*^*+/-*^ and C57 WT mice. **H**. KEGG pathway enrichment analysis for DEGs between C57-*En1*^*+/-*^ and C57 WT mice, including Parkinson’s disease. **I**. Heatmap showing genes included in the enriched pathways among DEGs between C57-*En1*^*+/-*^ and C57 WT mice. Red: up-regulated, blue: down-regulated. GO terms were separated according to their Ontology category (Biological Process, Cellular Compartment and Molecular Function). OF1: SwissOF1, C57: C57Bl/6J

We performed GO enrichment analysis on the DEGs identified between OF1-*En1*^+/-^ and OF1 WT to get further insight of enriched biological pathways along with their associated molecular functions and cellular localization. Actin filament organization (q value 0.002), semaphorin-plexin signalling involved in axon guidance (q value 0.002) and leukocyte migration (q value 0.004) were among the most enriched biological processes, all of which play key roles in the brain development by modulating axonal guidance and cell migration (Limoni and Niquille, 2021) (Fig. 3, C). The KEGG database did not identify enriched pathways, but Reactome pathway-based analysis identified DEGs to be enriched in pathways linked to axon guidance, nervous system development and semaphoring-dependent signalling, further reinforcing the GO enrichment results (Fig. 3, D). Genes involved in axonal cone growth and guidance, as well as nervous system development were downregulated in OF1-*En1*^+/-^ compared to OF1 WT mice. DEGs present in several of the enriched pathways include noteworthy candidates involved in neuronal development and survival, including *Cck, Anxa1, Nos1, Nos3*, Plxna2, *Plxna4, Plxnd1* and *Sema3a* (Fig. 3, E).

Despite the low number of DEGs in C57-*En1*^+/-^ vs C57 WT mice, GO enrichment analysis identified biological processes linked to ATP generation in mitochondria. For the cellular compartment category, the enrichment was related to the mitochondrial inner membrane and specifically the respiratory chain complex (Fig. 3, F). Reactome pathway analysis also identified pathways associated with energy metabolism in the mitochondria, including respiratory electron transport, ATP production and tricarboxylic acid cycle (Fig. 3, G). Interestingly, pathways related with reactive oxygens species (ROS) production, oxidative phosphorylation (OxPhos) and neurodegenerative diseases were enriched in C57-*En1*^+/-^ vs C57 WT mice according to KEGG (Fig. 3, H). Of note, genes related to OxPhos and ATP generation including *Ndufa1* (part of complex I), *mt-Cytb* (part of complex III), *mt-Co1, mt-Co2, mt-Co3* (constituents of complex IV) along with constituents of mitochondrial ribosomes (*Mrps21*) were all upregulated in C57-*En1*^+/-^ compared to C57 WT mice. It is also worth mentioning that, although the pathways were only enriched in OF1-*En1*^+/-^ vs OF1 WT, individual DEGs in C57-*En1*^+/-^ vs C57 WT are involved in extracellular matrix organisation (*Ext2, Krt77, Mtmr14*).

### 5.5. C57 mice display uniquely regulated genes related to the electron transport chain in mitochondria

To identify genes and pathways associated with strain-dependent susceptibility to PD-like pathology, we compared transcriptional profiles in SNpc of OF1 and C57 mouse strains at 1 week. Between the WT strains, there were 366 DEGs, 210 with higher and 156 with lower expression in OF1 WT compared to C57 WT mice. Between *En1*^+/-^ mice, there were 355 DEGs, 184 with higher and 151 with lower expression in OF1-*En1*^+/-^ compared to C57-*En1*^+/-^ mice (Fig. 4, A). Among the DEGs, 162 occurred in both the comparison between WT and between *En1*^+/-^ mice and were thus independent of *En1*^+/-^ (Fig. 4, B).

**Figure 4.**
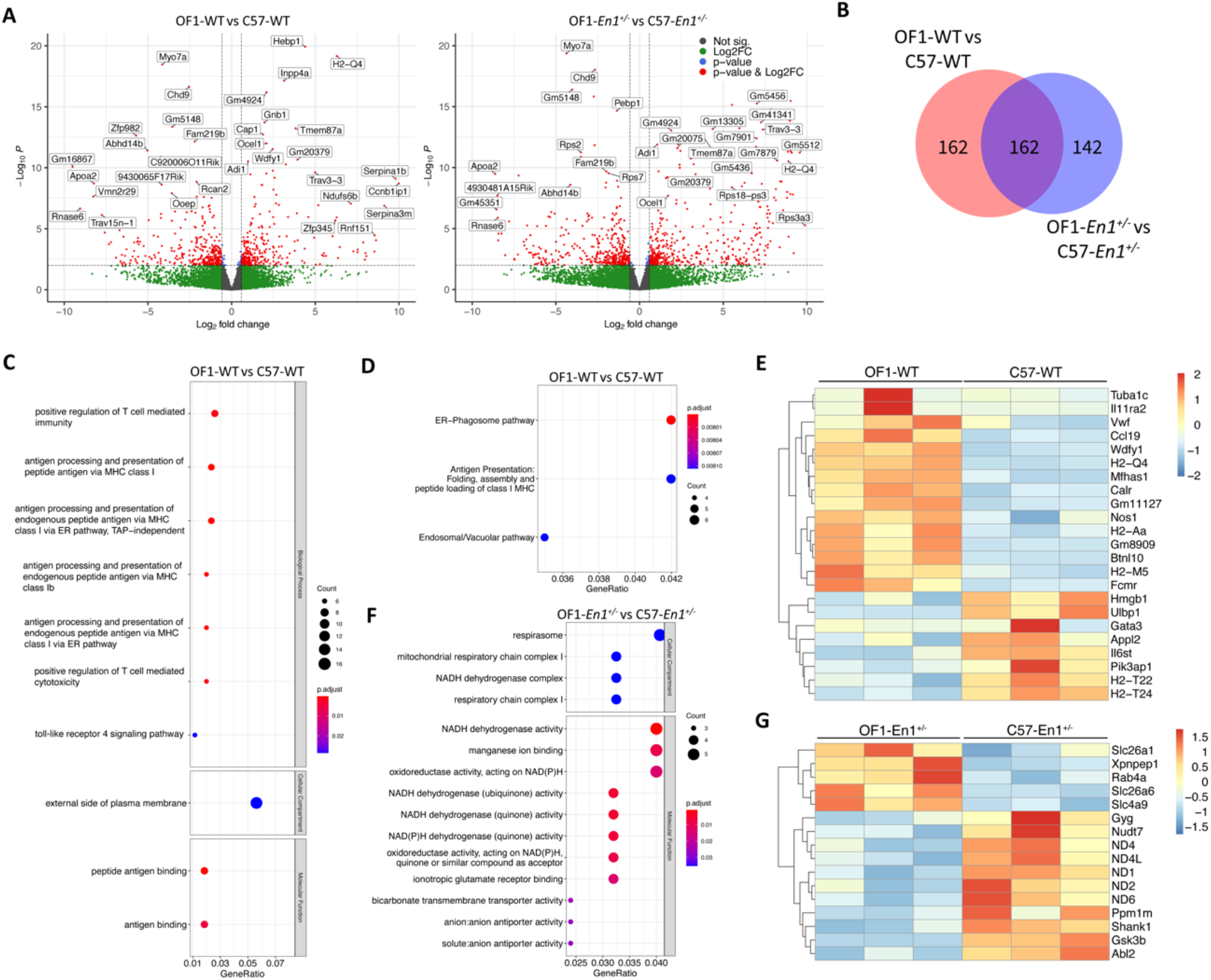
Analysis of transcriptomic differences between OF1 andC57 mice with wild-type (WT) genotype and with Engrailed 1 hemizygosity (*En1*^*+/-*^). **A**. Volcano plots with all expressed genes show transcriptomic differences in OF1 WT vs. C57 WT mice (left) and in OF1-*En1*^*+/-*^ vs. C57*-En1*^*+/-*^ mice (right). Vertical lines indicate fold change (FC) of 1.5. **B**. Venn diagram illustrating the overlap between differentially expressed genes (DEGs) detected in OF1 WT vs. C57 WT and in OF1*-En1*^*+/-*^ vs. C57*-En1*^*+/-*^ mice. **C**. Gene Ontology (GO) enrichment analysis of DEGs between OF1 WT and C57 WT mice. **D**. Reactome pathway analysis for DEGs between OF1 WT and C57 WT mice reveals pathways associated with vesicular trafficking and antigen presentation. **E**. Heatmap of genes included in the enriched pathways among DEGs between OF1 WT and C57 WT mice. OF1 WT mice show a higher expression of genes associated with antigen presentation and vesicle trafficking. Red: up-regulated, blue: down-regulated. **F**. GO enrichment analysis of the 142 DEGs exclusive to OF1-*En1*^*+/-*^ vs. C57-*En1*^*+/-*^ mice. The analysis shows enrichment in GO terms associated to energy metabolism in mitochondria. **G**. Heatmap of genes included in the enriched pathways among DEGs between OF1-*En1*^*+/-*^ and C57-*En1*^*+/-*^ mice. C57*-En1*^*+/-*^ mice consistently show higher expression of genes associated with mitochondrial respiration and oxidative phosphorylation. Red: up-regulated, blue: down-regulated. GO terms were separated according to their Ontology category (Biological Process, Cellular Compartment and Molecular Function). OF1: SwissOF1, C57: C57Bl/6J

The 366 DEGs in OF1 WT vs C57 WT were enriched in GO-terms such as antigen processing and presentation via major histocompatibility complex (MHC) class I, positive regulation of T cell mediated cytotoxicity and toll-like receptor 4 signalling pathway (Fig. 4, C). Reactome pathway analysis revealed that ER-phagosome and antigen presentation pathways were enriched, thus further supporting the GO enrichment results (Fig. 4, D). Among the genes present in the enriched pathways were genes encoding proteins involved in antigen presentation, including H2-Qa, H2-Aa, Gm11127, H2-T22 and H2-T24 (Fig. 4, E)

No enrichment was observed for the 355 DEGs in OF1-*En1*^+/-^ vs C57-*En1*^+/-^ when analysing with GO, KEGG or Reactome pathway enrichment analysis. However, the 142 DEGs that were exclusive to *En1*^+/-^, and not found in WT strain comparisons, were significantly enriched in molecular functions related to NADH dehydrogenase activity and ion transporter activity. No biological processes were significantly enriched; however, it is worth noting that enrichment for cellular compartment ontology showed again localization to the mitochondria, particularly in the respirasome (Fig. 4, F), similar to DEGs found in C57-*En1*^*+/-*^ vs C57 WT mice.

When assessing the genes associated with enriched molecular functions in respirasome, we observed that OF1-*En1*^+/-^ mice had a lower expression of OxPhos-related genes compared to C57-*En1*^+/-^ mice. GO terms like respirasome, mitochondrial respiratory chain complex I and NADH dehydrogenase activity included genes encoding constituents of complex I (mt-*Nd1, mt-Nd2, mt-Nd4, mt-Nd4L, mt-Nd6)* and complex V ATP synthase (mt-*Atp8*). This could indicate that C57-*En1*^+/-^ mice, but not OF1-*En1*^+/-^, compensate for impairments caused by *En1*^+/-^ with extra ATP production.

DEGs related to oxidative phosphorylation were found in both C57-*En1*^+/-^ vs C57-WT and OF1-*En1*^+/-^ vs C57-*En1*^+/-^. Given the importance of energy metabolism and mitochondrial functionality in neurodegenerative diseases we decided to see how this differential expression maps into the OxPhos chain and further look for differential expression of related genes between the two remaining comparisons, i.e., OF1-WT vs C57-WT and OF1-*En1*^+/-^ vs OF1-WT (Fig. 5). Despite not being among the enriched pathways, there was a significant higher expression of genes encoding complex I proteins in the OF1-WT strain when compared to the C57-WT strain (Fig. 5, B). Interestingly, when mapping the DEGs due to genetic background differences in the context of *En1* hemizygosity there were also differences in genes coding for complex I proteins (Fig. 5, C) with lower expression of 5 genes in OF1-*En1*^+/-^ mice compared to C57-*En1*^+/-^. However, there was no OxPhos-related differential gene expression when comparing OF1-*En1*^*+/-*^ to OF1-WT mice. Finally, when mapping the genes enriched in OxPhos pathways that were differentially expressed in C57-*En1*^+/-^ vs C57 WT mice, we observed higher expression of genes coding for constitutes of Complexes I (*Ndufa1*), III (mt-*Ctyb*) and IV (mt-*Cox1, mt-Cox2 and mt-Cox3*) (Fig. 5, D).

**Figure 5.**
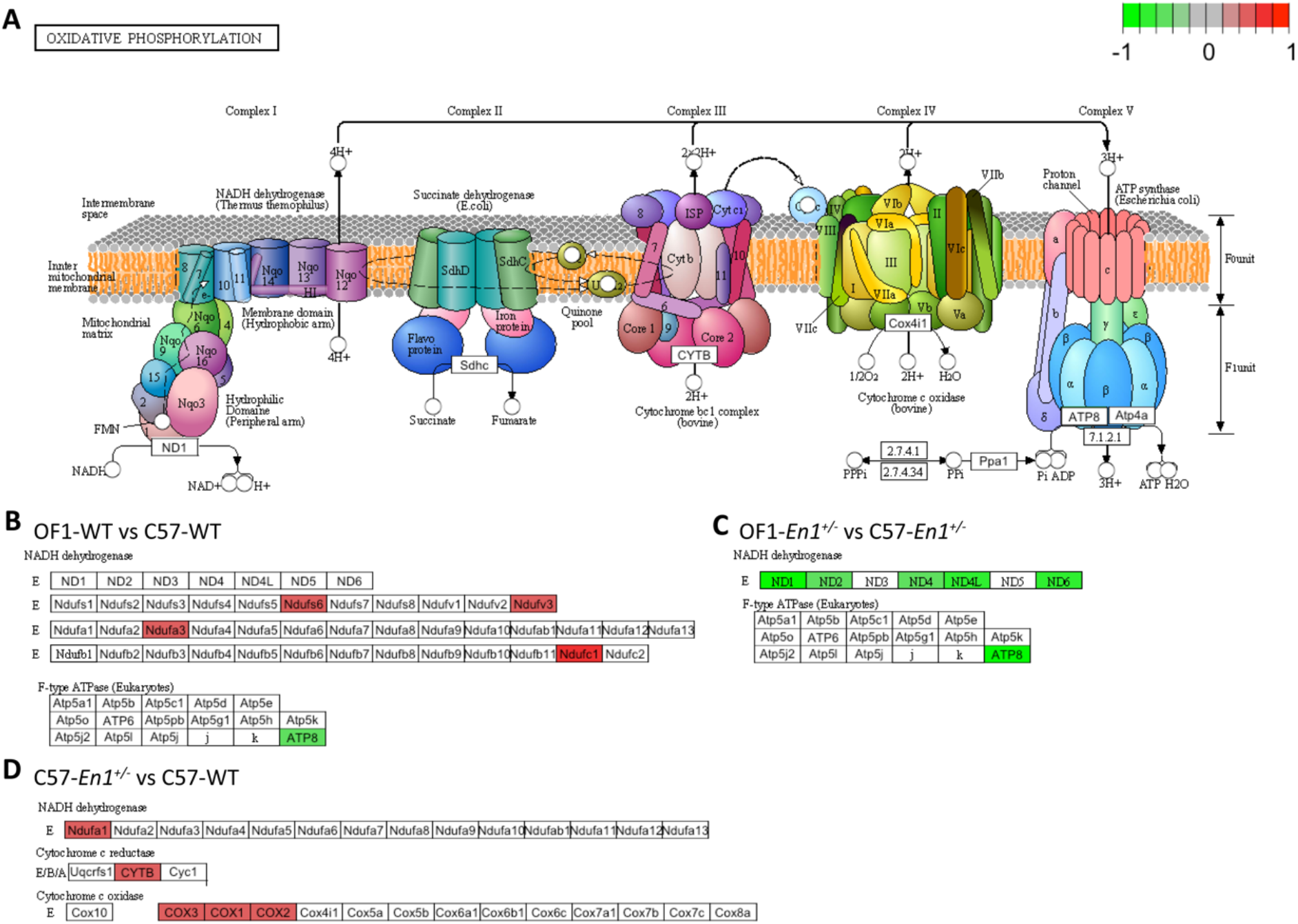
Genes with higher expression in C57 compared to OF1 mice encode components of the oxidative phosphorylation pathway. **A**. KEGGs representative diagram of the oxidative phosphorylation pathway. **B**. OF1 wild-type (WT) mice display higher expression of genes coding for constituent subunits of complex I and lower expression of ATP8 (in complex V) compared to C57 WT mice. **C**. In the context of En1 hemizygosity, OF1-*En1*^*+/-*^ mice have a lower expression of genes coding for complex I and V subunits compared to C57-*En1*^*+/-*^ mice. **D**. C57-*En1*^*+/-*^ mice have a higher expression of genes coding for components of complexes I, III and IV compared to C57 WT mice.

Overall, this data suggests that genetic background differences between the OF1 and the C57 strains have a direct effect on gene expression in the SNpc, particularly in pathways associated to energy metabolism in the mitochondria, and ultimately modulate the presence of PD-like pathology.

## 6. Discussion

The importance of the genetic landscape on PD incidence has been widely established by genetic studies, which have shed light on how different genetic variants affect susceptibility to the disease (Nalls et al., 2014, 2019; Deng et al., 2018; Puschmann et al., 2019; Yao et al., 2021; Brolin et al., 2022). However, unravelling genetic susceptibility in multifactorial diseases like PD is challenging due to its underlying complexity involving interactions of genetic and environmental factors (Bloem et al., 2021). In this context, identification of the cellular and physiological mechanisms that mediate genetically determined susceptibility is essential for the understanding the onset and progression of PD.

The *En1*^*+/-*^ mouse model has been shown to reproduce many of the characteristic traits of PD, including a strong genetic background component manifested as differential susceptibility to PD-like pathology in different mouse strains. Previous works have shown that OF1*-En1*^*+/-*^ mice exhibit a nigrostriatal neurodegeneration of nigrostriatal dopaminergic neurons (Sonnier et al., 2007; Nordström et al., 2015) while C57*-En1*^*+/-*^ mice lack this PD-like phenotype (Sgadò et al., 2006c). Furthermore, polymorphisms in the *En1* gene have been associated with idiopathic PD in a small cohort study (Haubenberger et al., 2011). Our previous genetic linkage analysis in a C57×OF1-*En1*^+/-^ F2 intercross identified multiple interacting QTLs linked to the PD-like phenotype induced by *En1* hemizygosity (Kurowska et al., 2016). The complex genetic regulation of the traits makes further QTL mapping difficult and call for complementary strategies. We therefore decided to use transcriptomic analyses to identify factors underlying the susceptibility to *En1*^*+/-*^-induced PD-like pathology. To our knowledge, this is the first work to perform a strain differential susceptibility study on the SNpc of the *En1*^+/-^ mice at a transcriptomic level.

Nigrostriatal axonal pathology has been proposed as an early sign of neurodegeneration in PD before evident loss of neuronal bodies in the SNpc (Burke and O’Malley, 2013; Kordower et al., 2013). Our results show that both OF1-*En1*^+/-^ and C57-*En1*^+/-^ mice exhibit early signs of neurodegeneration at 4 weeks in the form of nigrostriatal axonal swellings, but that progressive axonal pathology and loss of dopaminergic neurons cell bodies is only seen in OF1-*En1*^+/-^ mice. This confirms the presence of a genetic background-dependent susceptibility, consistent with previous findings (Sonnier et al., 2007). It is worth noting that at both 4 and 16 weeks, C57 WT and C57-*En1*^*+/-*^ mice have lower counts of dopaminergic neurons in the SNpc compared to the OF1 WT strain. This suggests that the observed resistance to the PD-like phenotype in C57-*En1*^*+/-*^ mice is not due to a neuronal reserve in the SNpc exerting a buffering effect, but rather due to other involved mechanisms. Instead of SNpc cell numbers, cellular function and morphology may be more important for motor function, and the Collaborative Crossing Consortia reported that low performance on motor test was associated with striatal axonal branching rather than TH^+^ area in the SNpc (Thomas et al., 2021).

The observed enrichment in OxPhos related terms, despite a low number of DEGs between C57-*En1*^*+/-*^ and C57 WT mice, suggests this process to have an important role in the resistance to nigrostriatal neurodegeneration observed in C57 mice. OxPhos and mitochondria have been shown to play a key role in DN degeneration, and this was reflected in PD being amongst the enriched KEGG pathways (Murali Mahadevan et al., 2021a; Connelly et al., 2023; Kalra, 2023). Furthermore, *Keap1* had a lower expression in C57-*En1*^+/-^ compared to C57 WT mice. The encoded Kelch-like ECH-associated protein 1 acts as suppressor of *Nrf2* which drives anti-inflammatory and antioxidative responses and is of high relevance to PD (Kopacz et al., 2020)(Lin and Beal, 2006). The lower Keap1 expression in C57-*En1*^+/-^ mice could thus disinhibit *Nrf2* and potentially have neuroprotective effects.

When analysing the effect of *En1*^+/-^ in the OF1 strain, DEGs were enriched in processes of nervous system development, including cell migration and axonal growth, which could reflect early changes associated with proper synapsis development. Particularly, semaphorin 3a (Sma3a) and plexins (*Plxna2* and *4*), which had a lower expression in OF1-*En1*^+/-^ compared to OF1 WT mice, have been showed to be essential for neuronal development and axonal elongation (Tamariz et al., 2010; Limoni and Niquille, 2021). Furthermore, transcriptomic studies in PD patients have reported an upregulation of the human orthologues to *Sma3a* and *Plxna4* (REF here!), and rare variants in *PLXNA4* have been linked to PD (Schulte et al., 2013).

When analysing the effect of genetic background on the transcriptome of WT OF1 and C57 mice, processes related to T-cell mediated immunity, antigen presentation and toll-like receptor 4 signalling were significantly enriched. However, no clear pattern of differential expression of pro- or anti-inflammatory molecules was observed in any strain and highly polymorphic immune-related genes such as those encoding MHC molecules could have impacted on the mapping of transcripts to genes and influence strain comparisons. In contrast, a clear enrichment in respirasome-related genes was present when evaluating DEGs exclusive to *En1* hemizygous mice, and these genes had a higher expression in C57*-En1*^+/-^ compared to OF1*-En1*^+/-^ mice. Most of these genes encode protein subunits of the mitochondrial respiratory chain complex I. No difference in OxPhos-related genes was found when comparing OF1-*En1*^*+/-*^ to *OF1* WT mice, suggesting that a neuroprotective effect mediated by higher expression of OxPhos-related genes is specific to C57*-En1*^*+/-*^ mice. Given that EN1 has been shown to protect mouse midbrain dopaminergic neurons against complex I insults (Alvarez-Fischer et al., 2011), this further reinforces that genetic components outside the *En1* locus could be driving ta complex I-mediated neuroprotective effect in C57 mice.

The consistent detection of differential gene expression of mitochondrial genes and genes that code for proteins in the OxPhos chain is promising given the relevance of this pathway in neurodegenerative processes. There is strong evidence linking mitochondrial impairment and energy metabolism to PD progression and pathogenesis in different model organisms (Briston and Hicks, 2018; Ordonez et al., 2018; Ikuno et al., 2021; Murali Mahadevan et al., 2021b). Our results suggest that early mechanisms, before the onset of neurodegeneration, takes place in the mitochondria and halts the neurodegenerative process in the C57-*En1*^+/-^ mice up to 16 weeks of age. Further experiments are needed to understand the exact mechanisms and the impact that these gene expression changes could have on nigral physiology.

Some considerations are needed when interpreting these results. We cannot rule out that the nigrostriatal pathology would evolve into dopaminergic neurons loss in the SNpc of C57-*En1*^+/-^ mice at an advanced age. Therefore, we cannot conclude if the neurodegenerative process is halted or delayed, but both scenarios are of high relevance to PD, where both halted and delayed disease progression would have significant clinical importance.

A limitation of this work is that bulk-RNAseq does not address the contribution of different cell types. Despite being able to obtain samples which are highly enriched in DN from the SNpc, we cannot rule out the possibility of contamination with neighbouring cells. Differences in sample cell composition have been shown to be a confounder in transcriptomic analyses performed in PD cohorts (Nido et al., 2020). However, in those cases, samples were dissected when the disease was advanced and therefore should lack many dopaminergic cells (Alves et al., 2009; Gaare et al., 2018).

The results from this study provide further evidence on the importance of genetic background for the onset and progression of nigrostriatal degeneration. The data provides valuable insight into how differential expression of genes encoding mitochondrial proteins before the onset of dopaminergic neurodegeneration is associated to differential vulnerability of nigral dopaminergic neurons to PD-like pathology. The fact that mitochondrial function and turnover are critical processes in PD is well known, but the findings presented here that normal genetic variation between strains alters expression of genes encoding mitochondrial proteins open for new therapeutic targets to prevent onset and/or progression of PD. As an example, metabolic mapping can provide accurate readouts on how energy substrates are consumed and identify precise differences in anaplerotic and mitochondrial driven energy generation (McKenna et al., 2012; Walls et al., 2014).

In conclusion, this study shows that neuronal metabolism and mitochondrial physiology are key components of PD-related neurodegeneration and suggests that early expression changes in genes encoding mitochondrial proteins have a key impact on disease onset and progression.

## Acknowledgments

This research was supported by MultiPark – a Strategic Research Area at Lund University and by grants from the Swedish Research Council (VR), Olle Engkvist’s foundation, Parkinsonfonden, Nilsson-Elhe Endowments from the Royal Physiographic Society of Lund, Lindhés advokatbyrå, and Sigurd & Elsa Golje’s memorial foundation (LB/MS). We would also like to thank Jenny G. Johansson from the MultiPark Next Generation Sequencing facility.

## Figures and references

**Supplementary Figure 1.**
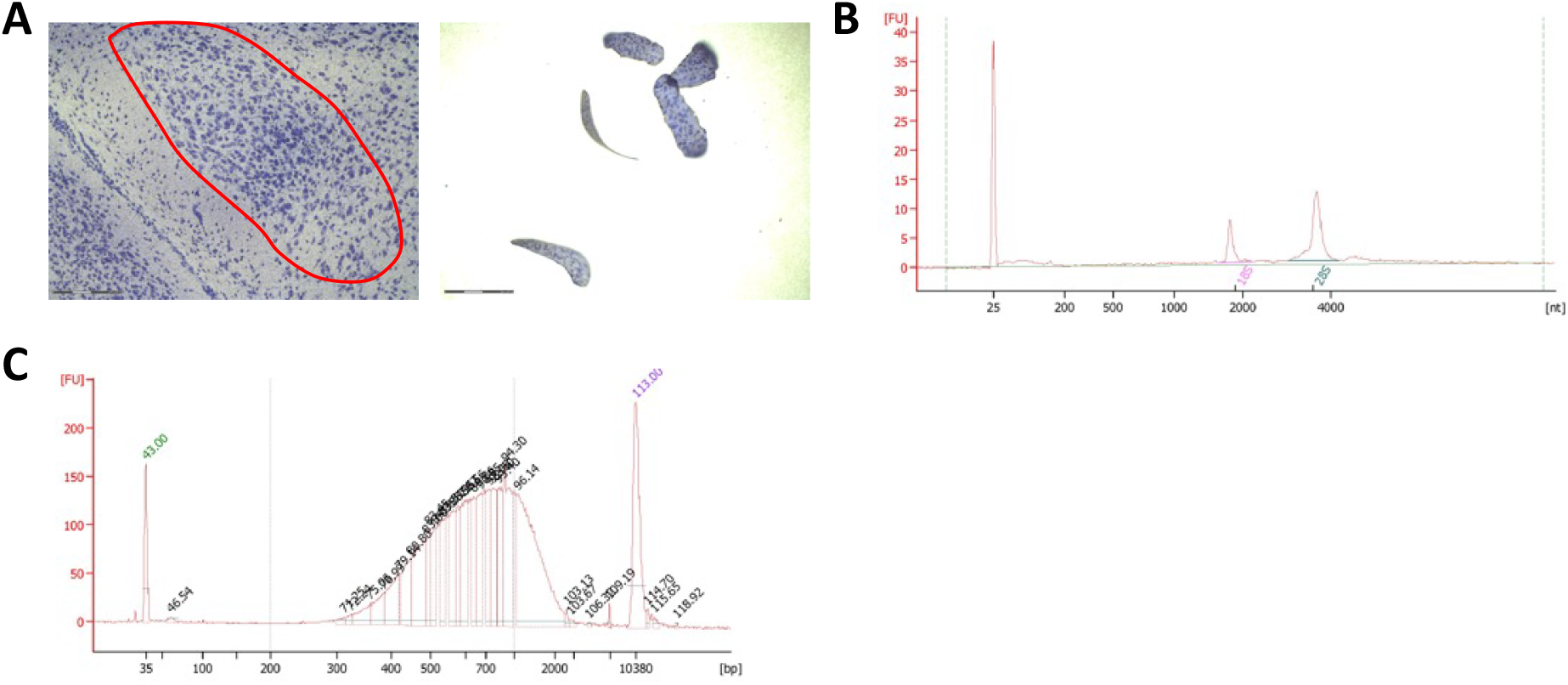
Laser capture microdissection (LCM) and RNA extraction. **A**. Representative images of substantia nigra pars compacta (SNpc) stained with cresyl violet prior to LCM. **B**. Representative electropherogram of intact RNA (based on the presence of the rRNA S18 and S28 peaks; RNA Integrity Number > 8), retrieved by direct lysis of microdissected tissue. **C**. cDNA libraries were successfully generated with the SMART-Seq®v4 Ultra®Low Input RNA Kit for Sequencing followed by shearing and sequencing library preparation.

**Supplementary Figure 2.**
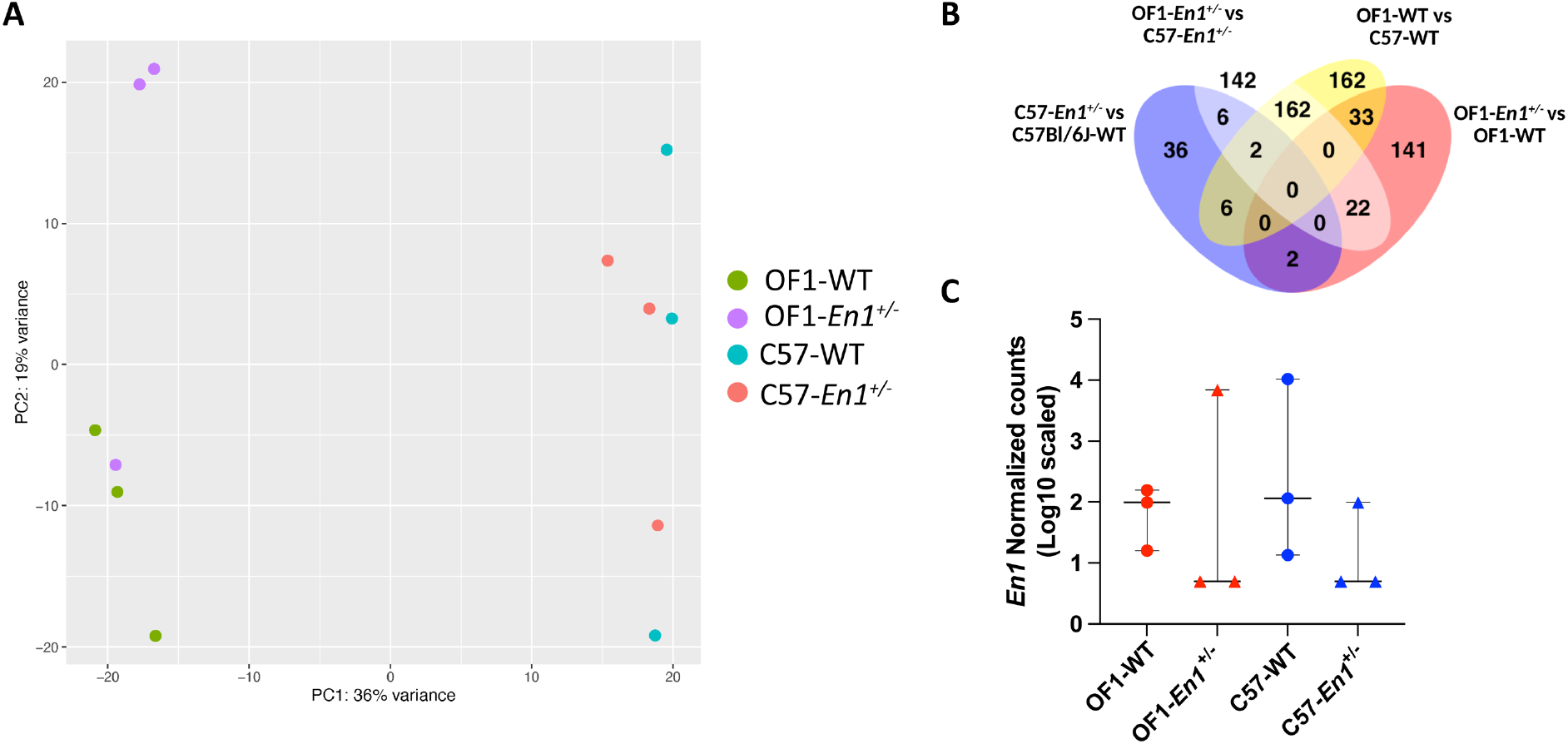
**A**. Principal component analysis (PCA) of the samples shows that the main effect is due to genetic background. **B**. Venn diagram illustrating the intersection between the four sets of differentially expressed genes (DEGs). **C**. Detected levels of *Engrailed 1* (*En1*) in the RNA-Seq data.

## Notes

**Conflict of interest** The authors declare no competing financial interests.

### Competing Interest Statement

The authors have declared no competing interest.

